# Bayesian Hidden Mark Interaction Model for Detecting Spatially Variable Genes in Imaging-Based Spatially Resolved Transcriptomics Data

**DOI:** 10.1101/2023.12.17.572071

**Authors:** Jie Yang, Xi Jiang, Kevin W. Jin, Sunyoung Shin, Qiwei Li

## Abstract

Recent technology breakthroughs in spatially resolved transcriptomics (SRT) have enabled the comprehensive molecular characterization of cells whilst preserving their spatial and gene expression contexts. One of the fundamental questions in analyzing SRT data is the identification of spatially variable genes whose expressions display spatially correlated patterns. Existing approaches are built upon either the Gaussian process-based model, which relies on *ad hoc* kernels, or the energy-based Ising model, which requires gene expression to be measured on a lattice grid. To overcome these potential limitations, we developed a generalized energy-based framework to model gene expression measured from imaging-based SRT platforms, accommodating the irregular spatial distribution of measured cells. Our Bayesian model applies a zero-inflated negative binomial mixture model to dichotomize the raw count data, reducing noise. Additionally, we incorporate a geostatistical mark interaction model with a generalized energy function, where the interaction parameter is used to identify the spatial pattern. Auxiliary variable MCMC algorithms were employed to sample from the posterior distribution with an intractable normalizing constant. We demonstrated the strength of our method on both simulated and real data. Our simulation study showed that our method captured various spatial patterns with high accuracy; moreover, analysis of a seqFISH dataset and a STARmap dataset established that our proposed method is able to identify genes with novel and strong spatial patterns.

## 1 INTRODUCTION

Recent advancements in spatially resolved transcriptomics (SRT) technology have fundamentally transformed our capacity to study cellular behavior at a molecular level, while preserving their spatial and gene expression contexts. This technological leap has opened new avenues for exploring complex biological systems at unprecedented levels of detail and accuracy. Efremova et al. (2020) and Liao et al. (2021) found that the positional context of gene expression is important to understanding tissue functionality and pathology changes, which highlights the pivotal role of SRT techniques. Broadly, SRT technologies are categorized into sequencing-based and imaging-based methods based on differences in RNA profiling: sequencing-based and imaging-based. Spatial transcriptomics, one of the next-generation sequencing (NGS) technologies, resolves gene expression profiles at a resolution of 100 *μm*. Spatial transcriptomics implemented by the 10x Visium platform achieved 55*μm* resolution, allowing for a detailed study of spatial organization. On the other hand, imaging-based technologies have revolutionized the field of transcriptomics by achieving single-cell resolution, with prominent examples such as sequential fluorescence in situ hybridization seqFISH (Ståhl et al., 2016), seqFISH+ (Eng et al., 2019), and multiplexed error-robust FISH (MERFISH) (Moffitt et al., 2018). Datasets profiled by SRT technologies have inspired the exploration of the spatial organization of gene expression within tissues. Cohorts with single-cell resolution motivate more biological analysis, such as cell-cell communication analysis *via* CellChat (Jin et al., 2021), characterization of ligand-receptor interactions between different cell types (Efremova et al., 2020) and so on. Hence, spatial information provided by imaing-based SRT data makes it more feasible to identify and quantify gene expression in specific regions of a tissue.

One of the most interesting questions arising along the development of SRT techniques is the identification of spatially variable genes (SVGs) whose expressions display spatially correlated patterns. Studies have found that SVGs demarcate clear spatial substructure, and are relevant to disease progression (Svensson et al., 2018; Hu et al., 2021). Various methods across different fields have been developed to identify SVGs, each capitalizing on distinct strengths. Trendsceek (Edsgärd et al., 2018) is built upon marked point processes to rank and evaluate the spatial pattern of each gene; however, it yields unsatisfactory performance (Sun et al., 2020) and is inhibited from scaling to large-scale data due to the expensive computational cost (Sun et al., 2020; Dries et al., 2021). SpatialDE (Edsgärd et al., 2018), SPARK (Sun et al., 2020), and BOOST-GP (Li et al., 2021) capture spatial correlation patterns by utilizing the Gaussian process. Specifically, SpatialDE models normalized gene expression levels via a multivariate Gaussian model with a spatial covariance function characterizing linear and periodic spatial patterns. SPARK models raw counts using a generalized linear spatial model with different periodic and Gaussian kernels. BOOST-GP models raw counts with a Bayesian zero-inflated negative binomial (ZINB) model with a squared exponential kernel covariance matrix. However, the performance of these kernel-based methods relies heavily on the resemblance between the underlying spatial expression patterns and the predefined kernel functions (Jiang et al., 2022). BinSpect (Dries et al., 2021), a non-model based method, identifies SVGs through statistical enrichment analysis of spatial network neighbors with binarized gene expression states. SpaGCN (Hu et al., 2021) defines SVGs as genes as those exhibiting differential expression among spatial domains and employs a deep learning model to identify these domains. BOOST-MI (Jiang et al., 2022) utilizes an energy-based modified Ising model to identify SVGs exclusively for sequencing-based SRT data, with the limitation that the spatial position of measured spots needs to be on the regular lattice grid. Compared to kernel-based models, energy-based interaction characterization enables the detection of broader types of gene spatial expression patterns.

As mentioned, gene expressions resolved by imaging-based SRT have single-cell resolution, which potentially unearths more biological insights. We aimed to develop a model that can identify SVGs with higher accuracy to be used on data from imaging-based SRT platforms, and uncover more biological mechanisms. Drawing inspiration from the success of energy-based models over kernel-based approaches (Jiang et al., 2022), we propose a novel joint Bayesian framework model, BOOST-HMI. This model utilizes a recently proposed energy function for mark interaction (Li et al., 2019). In particular, we adopt a ZINB mixture model to handle the unique data characteristics of SRT, including excess zeros and unknown mean-variance structures. Additionally, our method introduces a latent binary gene expression indicator to distinguish high and low expression states at the cellular level, thereby enhancing the model’s robustness against noise. Unlike BOOST-MI, our proposed BOOST-HMI is not constrained by the spatial distribution requirements of sequencing-based SRT data, making it versatile for imaging-based datasets where cells are randomly distributed. Furthermore, BOOST-HMI directly models raw counts within a joint Bayesian framework, addressing uncertainties associated with dichotomization. Our comprehensive simulation studies, covering various scenarios, demonstrate the superior accuracy of BOOST-HMI in detecting SVGs. We also applied our model to two real datasets: a mouse hippocampus seqFISH dataset and a mouse visual cortex STARmap dataset, where it successfully detected more spatial patterns and layer-specific SVGs, potentially unveiling novel biological insights.

The rest of the paper is organized as follows: section 2 introduces our ZINB mixture model for identifying SVGs from SRT count data and discusses the extension of the Bayesian mark interaction model to SRT data. In section 3, we describe the Markov chain Monte Carlo (MCMC) algorithms for posterior sampling and the resulting posterior inference. Finally, section 4 presents our method’s performance on simulated and real SRT datasets, compared with five other methodologies.

## 2 METHODS

In this section, we introduce a ZINB mixture model for directly modeling the imaging-based SRT count data, and a hidden mark interaction model to quantify the spatial dependency of latent binary gene expression levels. The schematic diagram of BOOST-HMI is shown in Figure 1.

**Figure 1.**
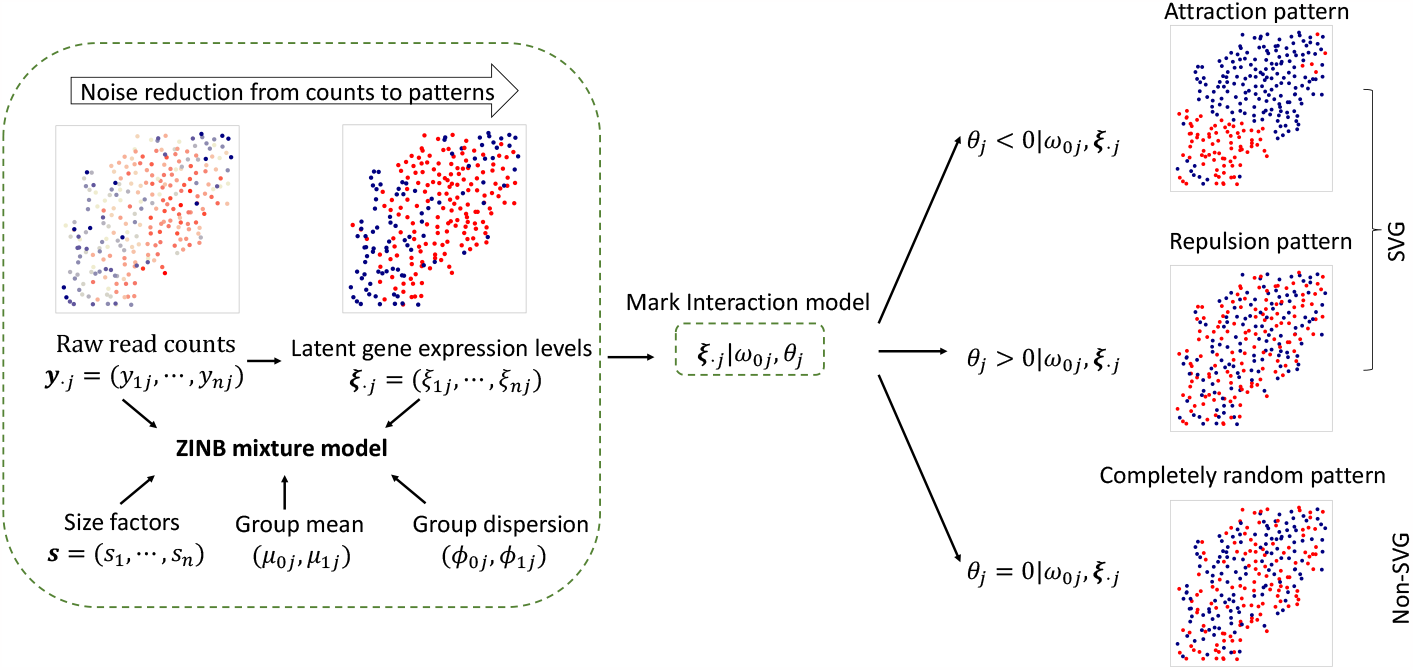
The schematic diagram of the proposed BOOST-HMI model.

Before introducing the models, we summarize the SRT data notations as follows. We denote the gene expression raw counts as a *n*-by-*p* matrix ***Y*** with each entry *y*_*ij*_ ∈ ℕ denoting the number of read counts for gene *j* at cell *i*. Every column ***y***_·*j*_ in ***Y*** denotes the expression counts across all measured cells for gene *j*, while each row ***y***_*i*·_ denotes the counts of all genes on cell *i* where *i* = 1, · · ·, *n, j* = 1, · · ·, *p*. As to geospatial profile, let a *n*-by-2 matrix *T* be the matrix for the spatial location of cells, where each row ***t***_*i*·_ = (*t*_*i*1_, *t*_*i*2_) ∈ ℝ ^2^ records the coordinates of cell *i* in the 2D Cartesian plane.

### 2.1 A ZINB mixture model for modeling gene expression count data

For the majority of SRT techniques, gene expression measurements obtained are in the form of counts. In the context of for imaging-based SRT platforms, gene expressions are collected as the count of barcoded mRNA corresponding to a particular transcript within a single cell (Zhao et al., 2022). Due to the characteristics of these measurements, observed count data often suffers from over-dispersion and zero-inflation. The negative binomial distribution can effectively account for the mean-variance relationship in the raw counts. Moveover, the gene expression count matrix ***Y*** is characterized by an inflated number of zeros, resulting from imaging sensitivity and hybridization efficiency (Zhao et al., 2022); therefore, we generalized the negative binomial (NB) model to the ZINB model to account for both the over-dispersion and the high sparsity level, i.e.,

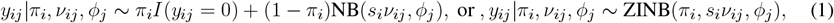

where parameter *π*_*i*_ ∈ [0, 1] represents the false zero proportion measured on cell *i*. NB(*ν, ϕ*) denotes a negative binomial distribution with mean *ν* and dispersion parameter *ϕ*. Consequently, the variance is *ν* + *ν*^2^*/ϕ*. 1*/ϕ* controls the overdispersion scaled by the square of mean *ν*^2^. The probability mass function is 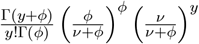. Given our particular circumstances, the NB mean is decomposed into two multiplicative effects, the size factor *s*_*i*_ and the expression level *ν*_*ij*_. The collection of ***s*** = (*s*_1_, · · ·, *s*_*n*_)^⊤^ reflects nuisance effects across cells. We follow SPARK Sun et al. (2020), setting *s*_*i*_ proportional to the summation of the total number of read counts across all genes for cell *i*, and combine it with a constraint of Π _*i*_ *s*_*i*_ = 1, which gives *s*_*i*_ = Σ_*j*_ *y*_*ij*_*/* Π _*i*_ Σ_*j*_ *y*_*ij*_. By setting the constraint for *s*_*i*_’s, we avoid the identifiability problem between *s*_*i*_’s and *ν*_*ij*_’s.

To denoise the relative expression levels, we aim to partition *ν*_*ij*_ into two groups by introducing the ZINB mixture model. Dichotomization has been widely applied as a step in the analysis of SRT data. BinSpect (Dries et al., 2021) and BOOST-MI (Jiang et al., 2022) discretize the normalized expression levels for each gene into two groups for more robust SVG detection results. Here, we introduce a latent binary gene expression level indicator vector ***ξ***_·*j*_ to denote the dichotomized expression profiles of each gene *j*. If *ξ*_*ij*_ = 1, gene *j* is highly expressed at cell *i*, and if *ξ*_*ij*_ = 0, gene *j* has low expression at cell *i*. A mixture model is suggested to allow different ZINB model parametrizations for high and low expression levels for gene *j*, in which we assume the raw expression count *y*_*ij*_ is generated one of two independent ZINB distributions with different means given the underlying binary indicator *ξ*_*ij*_,

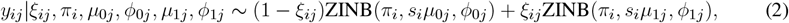

where *μ*_1*j*_ and *μ*_0*j*_, respectively, denote the group mean of read count for highly and lowly expressed genes. To guarantee that the mean expression level for a highly expressed gene is higher than a lowly expressed gene, we set a constraint for NB distribution mean across two expression levels: *μ*_1*j*_ *> μ*_0*j*_. *ϕ*_1*j*_ and *ϕ*_0*j*_ respectively represent the dispersion parameters of the NB model for the highly and lowly expressed gene.

To complete the model, we specify the following prior distributions: *μ*_0*j*_, *μ*_1*j*_ ∼ Ga(*a*_*μ*_, *b*_*μ*_), s.t. *μ*_1*j*_ *> μ*_0*j*_ *>* 0 and *ϕ*_0*j*_, *ϕ*_1*j*_ ∼ Ga(*a*_*ϕ*_, *b*_*ϕ*_). To create an environment conducive to model fitting, we introduce a latent variable *η*_*ij*_ to indicate whether a zero count *y*_*ij*_ is from the zero or NB component in Equation (1), and impose a Bernoulli prior *η*_*ij*_ ∼ Bern(*π*_*i*_), which can be further relaxed by formulating a Beta(*a*_*π*_, *b*_*π*_) hyperprior on *π*_*i*_, leading to a Beta-Bernoulli prior for *η*_*ij*_ with expectation *a*_*π*_*/*(*a*_*π*_ + *b*_*π*_).

### 2.2 A brief review of the Bayesian mark interaction model

Marked point interaction models are statistical models for spatial point pattern analysis with applications across diverse fields such as geostatistics, ecology, material physics, and so on (Edsgärd et al., 2018; Li et al., 2019). These models are designed to study the interactions among points with numerical or categorical marks in a planar region. Marked point models are receiving greater and greater focus in biology: for instance, Trendsceek applies the marked point process to identify SVGs (Edsgärd et al., 2018). The Bayesian mark interaction model, proposed by Li et al. (2019), is a full Bayesian model that characterizes spatial correlations among cell types from tumor pathology images.

Let (*t*_*i*1_, *t*_*i*2_) ∈ ℝ^2^, *i* = 1, …, *n* be the *x*- and *y*-coordinates of point *i*. Let *G* = (*V, E*) denote an interaction network with a finite set of points *V* and a set of direct interactions *E*. In the introduced Bayesian mark interaction model, we assume points have categorical marks. Here, we denote ***ξ*** = (*ξ*_1_, · · ·, *ξ*_*n*_)^⊤^ as the categorical marks of *n* points on the plane. *ξ*_*i*_ ∈ (1, · · ·, *Q*), *Q* ≥ 2 are the marks of point *i*.

The Bayesian mark interaction model formulates the pattern of marks ***ξ*** via the energy function, which is first introduced in statistical mechanics. The energy function has terms to account for both first- and second-order properties of the marked point data. Specifically, to model the interaction energy between two points, an exponential decay function with respect to the distance between the two points is used. Moreover, the Bayesian mark interaction model neglects interaction terms of point pairs from *E* when the corresponding distance is beyond a threshold *c*. Consequently, the model focuses on a sparse network *G*^*′*^ = (*V, E*^*′*^), where *E*^*′*^ includes edges joining pairs of points *i* and *i*^*′*^ with distance *d*_*ii*_*′ < c*. The setting of the distance threshold is added to avoid the high computation cost incurred when summing over *n* data points and (*n* − 1)*n/*2 interacting pairs of points with large *n*. Then, the potential energy of *G*^*′*^ is measured by two addictive terms,

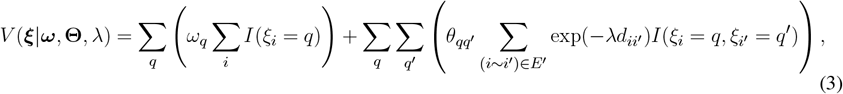

where *q, q*^*′*^ ∈ *{*1, · · ·, *Q}* are the categories of marks. ***ω*** = (*ω*_1_, · · ·, *ω*_*Q*_)^⊤^ and **Θ** = [*θ*_*qq*_*′*]_*Q×Q*_ are defined as first- and second-order intensities. (*i* ∼ *i*^*′*^) denotes the collection of interacting pairs of cells in 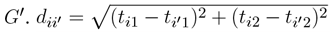 denotes the Euclidean distance between point *i* and *i*^*′*^. *λ* is the decay parameter of the distance between two points in the exponential decay function, where a larger *λ* makes energy diminish quickly with respect to the increase in point pair distance.

By restricting the interaction effect within radius *c*_*d*_, the Bayesian mark interaction model defines a local energy. According to the fundamental Hammersley-Clifford theorem (Clifford, 1990), a probability measure with a Markov property exists if we have a locally defined energy, called a Gibbs measure. This measure gives the probability of observing categorical marks associated with their locations in a particular state. We can write the joint probability on marks ***ξ*** as,

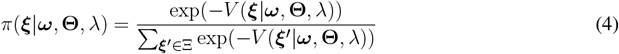

which is proportional to the exponential of the negative energy of marks ***ξ*** calculated by Equation (3). The denominator is a normalizing constant that needs to sum over the entire space **Ξ** of marks combination consisting of *Q*^*n*^ states, which is intractable even for a small size model.

The joint probability (Equation (4)) can be considered as the full data likelihood. To interpret the parameters clearly, we write the probability of observing point *i* having mark category *q* conditional on its neighborhood configuration,

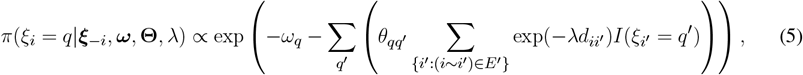

where ***ξ***_−*i*_ denotes the collection of all marks, except the *i*th one. Equation (5) shows that the probability of point *i* with mark *q* depends on parameter *ω*_*q*_, *θ*_*qq*_*′*, and the decay parameter *λ*. Parameters in Equation (5) are interpreted below. Suppose there is no interaction between any two points in the space, i.e., *θ*_*qq*_*′* = 0; then, the conditional probability of point *i* with *ξ*_*i*_ = *q* is *π*(*ξ*_*i*_ = *q*|·) ∝ exp(−*ω*_*q*_). Therefore, the model parameter *ω*_*q*_ is related to the abundance of points with mark *q*. Fixing ***ω*** as equal values, we obtain the conditional probability of point *i* with *ξ*_*i*_ = *q* is *π*(*ξ*_*i*_ = *q*|·) ∝ exp(−Σ_*q*_*′* [Σ_*{i*_*′*_:(*i*∼*i*_*′*_)∈*E*_*′*_*}*_ exp(−*λd*_*ii*_*′*) *I*(*ξ*_*i*_*′* = *q*^*′*^)]). The second-order intensity *θ*_*qq*_*′* quantifies the dependency of mark *q* with the nearby points, with mark *q*^*′*^ scaling by the distance decay function. A detailed parameter interpretation is provided by Li et al. (2019).

### 2.3 A hidden mark interaction model for identifying SVGs

In Section 2.1, we describe how our ZINB mixture model is used to convert read counts ***y***_·*j*_ for each gene *j* into their corresponding hidden binary states ***ξ***_·*j*_. This dichotomization process allows us to represent gene expression in a binary format. We then treat the spatial distribution of cells as a two-dimensional point process, with the binary gene expressions ***ξ***_·*j*_ serving as the markers of these points. This setup enables us to effectively employ the mark interaction model to assess the spatial correlations among these markers. For simplicity, as outlined in Section 2.2, we treat the decay parameter *λ* as a pre-defined hyperparameter *λ*_0_. Within this framework, the energy function can be expressed as follows:

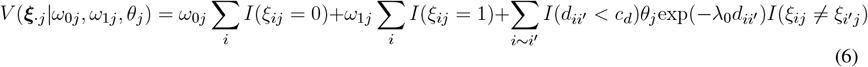

To interpret the model parameters, we provide the conditional probability of observing a high-expression level *ξ*_*ij*_ = 1 of gene *j* at cell *i*, given the expression levels of other cells for gene *j*:

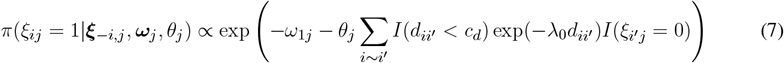

As introduced in Section 2.2, the model parameters ***ω***_*j*_ = (*ω*_0*j*_, *ω*_1*j*_)^⊤^ represent the first-order property, and *θ*_*j*_ reflects the second-order property of the spatial distribution of marks. Specifically, model parameters *ω*_0*j*_ and *ω*_1*j*_ are related to the abundance of cells with low and high expression levels of gene *j*, respectively. From Equation (7), if we assume *θ*_*j*_ = 0, then the distribution of *ξ*_*ij*_ for cell *i* is independent of the values of other cells and the proportion of high expression level *π*(*ξ*_*ij*_ = 1|·) = exp(*ω*_1*j*_)*/*(exp(*ω*_0*j*_) + exp(*ω*_1*j*_)). Assuming ***ω***_*j*_ are fixed, if the interaction parameter *θ*_*j*_ between lowly and highly expressed cells is positive, a larger probability of cell *i* belonging to the highly expressed group corresponds to fewer lowly expressed cells in the surrounding area (i.e., fewer terms *I*(*ξ*_*i*_*′*_*j*_ = 0)). In other words, if *θ*_*j*_ *>* 0, the gene expression level at cell *i* tends to be concordant with the majority of its neighboring cells, resulting in a repulsion pattern. If the interaction *θ*_*j*_ is negative, we can further infer that there is a spatial pattern in which cells with different expression levels are clustered, resulting in an attraction pattern. Consequently, we identified the association between cell interactions, which potentially results in a spatial pattern, and the second-order intensity parameter *θ*_*j*_. Parameter *λ*_0_ controls the change of interaction strength between a pair of points with respect to their distance. A larger *λ*_0_ causes the interaction between two cells to diminish faster, resulting in a smaller interactive neighborhood for each cell. As a hyperparameter, *λ*_0_ needs to be set appropriately to reflect the interaction neighborhood for cells.

An identifiability problem arises when adding a nonzero constant to *ω*_0*j*_ and *ω*_1*j*_, as it causes the joint probability *π*(***ξ***_·*j*_|·) to remain invariant. Therefore, we constrain *ω*_1*j*_ = 1 and establish prior distributions for *ω*_0*j*_ and *θ*_*j*_ to complete the parameter model settings for the hidden mark interaction model: 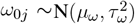 and 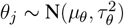.

## 3 MODEL FITTING

In this section, we introduce the MCMC algorithm for model fitting and posterior inference. Our model space consists of (***M***, **Φ, *H*, Ξ, *ω***_0_, ***θ***) with the underlying grouped gene expression levels ***M*** = [*μ*_*kj*_]_2*×p*_, the dispersion parameters **Φ** = [*ϕ*_*kj*_]_2*×p*_, the extra zero indicators ***H*** = [*η*_*ij*_]_*n×p*_, the binary expression level indicators **Ξ** = [*ξ*_*ij*_]_*n×p*_, the first-order intensity parameter ***ω***_0_ = (*ω*_01_, …, *ω*_0*p*_)^⊤^ and the interaction parameter ***θ*** = (*θ*_1_, …, *θ*_*p*_)^⊤^ in the mark interaction model. Each gene is examined independently by BOOST-HMI. We give the full posterior distribution for gene *j* as,

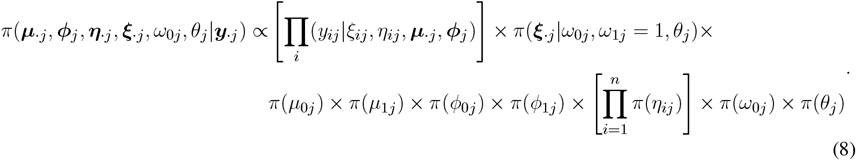

Our primary aim was to infer *ω*_0*j*_, *θ*_*j*_ and ***ξ***_·*j*_, which define the Gibbs probability measure based on the local energy function. We provide estimation and inference on first-order intensity *ω*_0*j*_, which represents the abundance of lowly expressed levels of gene *j*, and the second-order intensity *θ*_*j*_, which captures the spatial correlation between two expression levels. The estimated latent gene expression level indicator provides a robust estimation of the spatial organization of marks.

### 3.1 MCMC algorithms

We estimate *μ*_0*j*_, *μ*_1*j*_, *ϕ*_0*j*_ and *ϕ*_1*j*_ using the random walk Metropolis-Hastings (RWMH) algorithm. ***η***_·*j*_ and ***ξ***_·*j*_ are estimated with a Gibbs sampler. The Gibbs probability measure for the distribution of latent gene expression indicator ***ξ***_·*j*_ in Equation (7) omits an intractable normalizing constant 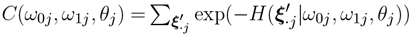, which makes the Metropolis-Hastings algorithm infeasible. For instance, to model a gene expression profile with *n* = 257 cells, we need to traverse 2^257^ ≈ 2.3 *×* 10^77^ different arrangements of ***ξ*** for every gene, which is a heavy computational burden. To overcome this issue, we use the double Metropolis-Hastings (DMH) algorithm proposed by Liang et al. Liang (2010) to estimate *ω*_0*j*_ and *θ*_*j*_ by canceling the intractable normalizing constant. The DMH is an efficient auxiliary variable MCMC algorithm. In contrast to other auxiliary MCMC algorithms, it does not require drawing the auxiliary variables from a perfect sampler, which usually increases computational cost (Møller et al., 2006). The full details of MCMC algorithms is described in the supplementary materials.

### 3.2 Posterior inference

Posterior inference of parameters *μ*_0*j*_, *μ*_1*j*_, *ϕ*_0*j*_, *ϕ*_1*j*_, *ω*_0*j*_, and *θ*_*j*_ is obtained by averaging the MCMC posterior samples after burn-in. We are interested in identifying the SVGs by summarizing the interaction parameter ***θ***. As stated in Section 2.3, investigating whether *θ*_*j*_ is positive or negative is of great importance to inferring the spatial expression pattern of gene *j*. To check the sign of *θ*_*j*_, we applied hypothesis testing ℳ _0_ : *θ*_*j*_ ≥ 0 versus ℳ _1_ : *θ*_*j*_ ≤ 0. If there is strong evidence to reject the null hypothesis ℳ _0_, we conclude that gene *j* is an SVG. The Bayes factor (BF) is computed to infer whether *θ*_*j*_ is positive or negative with statistical significance from the MCMC algorithm results. The Bayes factor measures the favorability of ℳ _1_ as

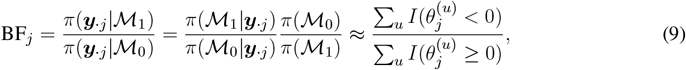

where *u* index the iteration and *U* is the total number of iterations after burn-in. The larger the BF_*j*_, the more likely gene *j* is an SVG with an attraction pattern. The smaller the BF_*j*_, the more likely gene *j* is an SVG with a repulsion pattern.

Another important parameter in our model is the latent gene expression level indicator ***ξ***_·*j*_. We summarize the posterior distribution of ***ξ***_·*j*_ via *maximum-a-posteriori* (MAP) estimates, which is the mode of the posterior distribution. A more comprehensive summary of ***ξ***_·*j*_’s is based on their marginal posterior probabilities of inclusion (PPI), where 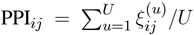. Then, the latent expression indicator indicates a high expression spot when PPIs are greater than a threshold *c*_*p*_:

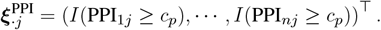

## 4 RESULTS

### 4.1 Simulation Study

We generated simulated data to evaluate the ability of BOOST-HMI to identify SVGs and provide a comparison with four competing methods: SpatialDE (Svensson et al., 2018), SPARK (Sun et al., 2020), BinSpect-kmean (Dries et al., 2021), and BinSpect-rank (Dries et al., 2021).

Spatial locations of simulated data were from the geospatial profile of the mouse hippocampus dataset field 43 (Shah et al., 2016) with *n* = 257 cells, which we present in Section 4.2. To generate the expression counts for gene *j*, the latent gene expression level indicators ***ξ***_·*j*_’s were first generated based on Equation (7) with three different values of *ω*_0*j*_ ∈ *{*1.4, 1, 0.6*}* and the fixed value of *ω*_1*j*_ = 1. These three values of ***ω***_*j*_ correspond to approximately 60%, 50%, and 40% lowly-expressed cells in ***ξ***_·*j*_. Additionally, we set four different values of *θ*_*j*_ ∈ *{*−2.5, −1.2, 1.9, 3.2*}* to generate SVGs with various patterns of attraction or repulsion. These values correspond to strong attraction, weak attraction, weak repulsion, and strong repulsion patterns, respectively. For the non-SVG, *θ*_*j*_ = 0 which indicates complete randomness and no spatial correlation. The distance threshold *c*_*d*_ and decay parameter *λ*_0_ were set as 0.15 and 20, respectively. We then simulated gene expression data from a ZINB model with three different group-mean ratios *r* ∈ *{*2, 5, 10*}* between high and low expression:

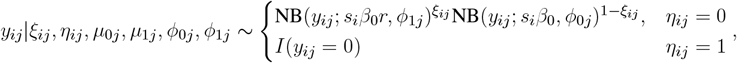

where the underlying baseline expression levels *β*_0_ = 10. In the simulation study, size factors ***s*** = (*s*_1_, · · ·, *s*_*n*_)^⊤^ were generated from log-N(0, 0.2^2^), and dispersion parameters *ϕ*_0*j*_, *ϕ*_1*j*_ in the NB model are generated from an exponential distribution Exp(1*/*10). Further, to imitate high sparsity and account for medium sparsity in real SRT data, we created three sets of sparsity levels, 0-10%, 10-20%, and 30-40%, and generated extra zero parameters ***η***_·*j*_’s correspondingly. Extra zeros were randomly selected and imputed into the generated gene expression count data. Thus, we considered three group-mean ratios and three sparsity levels, which is 3 *×* 3 = 9 scenarios in total. For each scenario, we simulated 30 replicates with *p* = 100 genes in each replicate, 10 out of which were SVGs.

Before estimating the parameters using BOOST-HMI, we specified the prior distributions. Noninformative gamma priors were specified for *μ*_0*j*_, *μ*_1*j*_, *ϕ*_0*j*_ and *ϕ*_1*j*_, i.e., *μ*_0*j*_ ∼ Ga(*a*_*μ*_, *b*_*μ*_) and *ϕ*_0*j*_ ∼ Ga(*a*_*ϕ*_, *b*_*ϕ*_). We set *a*_*μ*_, *b*_*μ*_, *a*_*ϕ*_, and *b*_*ϕ*_ to 0.01, which produced a gamma distribution with mean 1 and variance 100. Priors for *ω*_0*j*_ and *θ*_*j*_ were set to control the gene expression abundance and gene expression pattern, 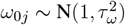 and 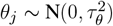. In the simulation study and real data analysis, we set *τ*_*ω*_ = 0.5, where the prior distribution of *ω*_0*j*_ indicates that the latent proportion of low expression cells ranges from 1% to 100% with a probability of 95%. *τ*_*θ*_ was set to 3.5 such that the prior for *θ*_*j*_ guarantees that *θ*_*j*_ falls within −6 to 6 with a probability of 92%. For hyperparameters in the energy function, we set the distance threshold *c*_*d*_ = 0.15 and expected the relationship of decay parameter *λ*_0_ and *c*_*d*_ to be exp(−*λ*_0_*c*_*d*_) = 0.05, which specifies the range of exponential decay function exp(−*λ*_0_*d*_*ii*_*′*) to be [0.05, 1] as *c*_*d*_ ≥ *d*_*ii*_*′* ≥ 0; therefore, we set *λ*_0_ as 20 correspondingly. As for the setting of the MCMC algorithm, we implemented BOOST-HMI in a gene-wise fashion. For each gene, we initialized model parameters by randomly drawing from their prior distributions. The MCMC algorithm is iterated *U* = 10, 000 times after 10, 000-iteration burn-in. The algorithm was implemented in `R` and `Rcpp`. As mentioned in Section 3.2, BOOST-HMI identifies SVGs based on Bayes Factors (BFs). In our study, we select a BF threshold of 10, which indicates strong evidence in favor of the ℳ_1_ (Kass and Raftery, 1995).

We implemented the other four competing methods with their default settings. SpatialDE, BinSpect and SPARK use *P* -values to select SVGs. To control type-I error rate, the Benjamini-Hochberg (BH) (Benjamini and Hochberg, 1995) procedure was used to adjust *P* -values from SpatialDE and BinSpect. We specifically avoided adjusting *P* -values from SPARK since its raw *P* -values are calibrated by the Cauchy combination rule (Sun et al., 2020; Liu et al., 2019). For *P* -values, the threshold was set to 0.05.

Our task is to evaluate the ability of each method to correctly identify underlying SVGs from the simulated dataset, which can be defined as a binary classification problem; therefore, to evaluate the performance of the five methods, we employed two performance metrics for binary classification problems: First, we used the area under the curve (AUC) (Bradley, 1997) of the receiver operating characteristic (ROC) (Fukunaga, 2013). The ROC is a plot of the true positive rate against the false positive rate for different classification thresholds. The AUC is a single value ranging from 0 to 1, with a higher value indicating better classification performance.

Figure 2 displays a boxplot of AUCs calculated by the aforementioned five methods over 30 replicates across nine scenarios. It clearly suggests that BOOST-HMI achieves superior performance compared to the other methods, especially for when there was high sparsity. BinSpect-kmeans and BinSpect-rank showed competitive performance when there was no zero-inflation, i.e., when the sparsity level was between 0% to 10%, regardless of the different group ratios; however, these methods showed decreasing AUCs as sparsity level increased. SPARK and SpatialDE suffered from a limited ability to detect SVGs from low expression variability or high zero-inflation scenarios. Between SPARK and SpatialDE, the simulation study showed that SPARK has better SVG detection power over SpatialDE, which is consistent with the conclusion drawn by Sun et al. (2020) and Jiang et al. (2022). In summary, BOOST-HMI achieved satisfactory performance and is robust against different group expression level ratios and sparsity levels.

**Figure 2.**
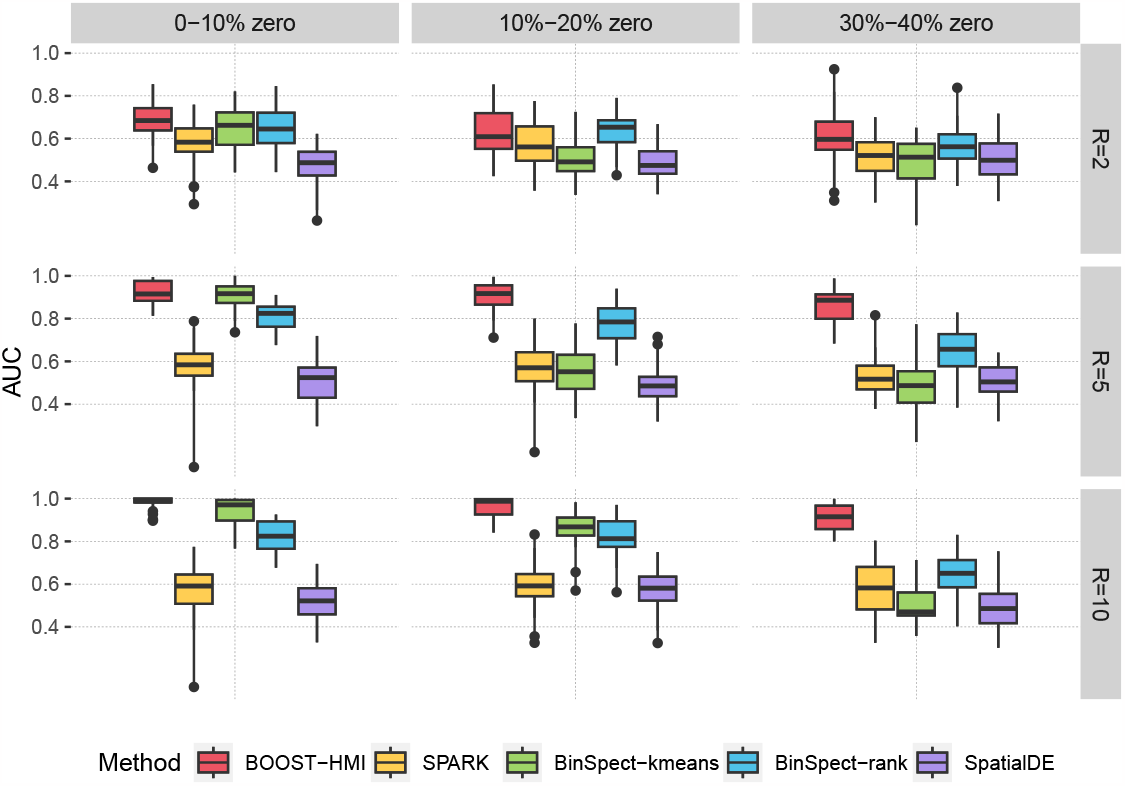
Simulation study: The boxplots of AUCs achieved by BOOST-HMI, SPARK, SpatialDE, BinSpect-kmeans, and BinSpect-rank across nine scenarios.

The second metric we used is the Matthews correlation coefficient (MCC). MCC is a summary value that examines the binary classification performance under a specific cutoff, i.e., BF or *P*−value thresholds for our study. It has values ranging from −1 to 1, incorporating true positives, true negatives, false positives and false negatives. A larger MCC value, such as 1, corresponds to an excellent classifier, while a negative MCC indicates a strong disagreement between prediction and observation. Table 1 summarizes the average MCCs obtained in the simulation study across the five methods. The result is consistent with our conclusion from our analysis of the AUC: BOOST-HMI achieved the highest power under high zero-inflation, while all other four methods suffered from the high number of false zeros. In the scenario without zero inflation, BinSpect-kmeans stood out, and BinSpect-rank, SPARK, and BOOST-HMI showed competitive performance in identifying SVGs.

**Table 1.** Simulation study: The averaged MCCs, with standard deviations in parentheses, achieved by BOOST-HMI, BinSpect-kmeans, BinSpect-rank, SPARK, and SpatialDE across nine scenarios.

|  | 0 - 10% zeros |  |  |
| --- | --- | --- | --- |
| | $R = 2$ | $R = 5$ | $R = 10$ |
| BinSpect-kmeans | 0.182 (0.159) | 0.653 (0.118) | 0.714 (0.085) |
| BinSpect-rank | 0.199 (0.193) | 0.476 (0.123) | 0.488 (0.139) |
| SPARK | 0.255 (0.162) | 0.502 (0.164) | 0.524 (0.153) |
| SpatialDE | 0.009 (0.111) | −0.009 (0.063) | 0.027 (0.109) |
| BOOST-HMI | 0.231 (0.141) | 0.449 (0.084) | 0.510 (0.063) |
|  | 10% - 20% zeros |  |  |
| | $R = 2$ | $R = 5$ | $R = 10$ |
| BinSpect-kmeans | 0.007 (0.092) | 0.121 (0.134) | 0.601 (0.142) |
| BinSpect-rank | 0.173 (0.166) | 0.482 (0.120) | 0.537 (0.153) |
| SPARK | 0.113 (0.146) | 0.271 (0.153) | 0.373 (0.123) |
| SpatialDE | −0.039 (0.043) | 0.005 (0.093) | 0.031 (0.120) |
| BOOST-HMI | 0.160 (0.176) | 0.433 (0.091) | 0.473 (0.065) |
|  | 30% - 40% zeros |  |  |
| | $R = 2$ | $R = 5$ | $R = 10$ |
| BinSpect-kmeans | 0.002 (0.086) | 0.005 (0.083) | −0.006 (0.105) |
| BinSpect-rank | 0.060 (0.139) | 0.243 (0.157) | 0.226 (0.142) |
| SPARK | 0.014 (0.103) | 0.133 (0.150) | 0.144 (0.155) |
| SpatialDE | −0.016 (0.071) | −0.001 (0.097) | 0.016 (0.110) |
| BOOST-HMI | 0.098 (0.158) | 0.406 (0.088) | 0.427 (0.088) |

### 4.2 Application to the mouse hippocampus seqFISH data

The mouse hippocampus dataset is a public seqFISH dataset with 21 field replicates collected on a third coronal section (Shah et al., 2016). Following SpatialDE and SPARK protocols, we analyzed the field 43 dataset, which contains *p* = 249 genes measured on 257 cells with spatial location preserved. Out of 249 genes, 214 were selected from a list of transcription factors and signaling pathway components, and the remaining 35 were selected from cell identity markers. Quality control was performed following SPARK protocol (Sun et al., 2020) and the original study. We filtered out cells with *x*-or *y*-axis values exceeding 203-822 pixels to tackle border artifacts. After filtering, *n* = 131 cells were included for the following analysis. We excluded SpatialDE due to its unsatisfactory performance in SVG detection in the simulation study. Prior settings, MCMC algorithm implementation, and significance criteria were identical to what was described in Section 4.1. Five independent MCMC chains were sampled and diagnosed for algorithm convergence. Algorithm convergence was checked based on the BF vector. BFs from five chains were highly correlated with Pearson correlation coefficients ranging from 0.90 to 0.99. We further checked algorithm convergence using the potential scale reduction factor (PSRF) (Gelman and Rubin, 1992; Brooks and Gelman, 1998) on posterior samples of *θ*_*j*_’s and *ω*_0*j*_’s. If multiple chains converge to the target posterior distribution, the PSRF will be close to one. In our analysis, the PSRFs were below 1.2, suggesting convergence of the MCMC algorithms. Posterior samples obtained from the quintuplet of MCMC chains were amalgamated for subsequent analysis.

Among the *p* = 249 genes analyzed in the mouse hippocampus dataset, SPARK identified 17 SVGs, while BOOST-HMI detected 22 SVGs. Notably, BOOST-HMI successfully detected 16 cell identity markers previously presented by Shah et al. (2016), whereas SPARK identified 14 markers. In comparison, BinSpect-kmeans and BinSpect-rank were more aggressive, respectively identifying 38 and 44 SVGs. A Venn diagram in Figure 3(D) showcases the overlap of SVGs identified by the four methods. Among them, BOOST-HMI and SPARK shared 12 SVGs in common. Only one SVG detected by BinSpect-kmeans overlapped with that from BOOST-HMI, and none overlapped with that from SPARK. None of the SVGs detected by BinSpect-rank were detected by either SPARK or BOOST-HMI. We further visualized the spatial pattern for each SVG using the marginal PPI of the posterior samples of the hidden gene expression indicator ***ξ***_·*j*_. Figure 3(A) displays the spatial patterns of SVGs detected by SPARK and BOOST-HMI, while Figures S1 and S2 in the supplementary materials depict the spatial patterns for SVGs detected by BinSpect-kmeans and BinSpect-rank, respectively. Among the 12 common SVGs identified by SPARK and BOOST-HMI, strong spatial repulsion patterns between high- and low expression genes are evident across 131 cells. Notably, a larger Bayes factor (shown in parentheses) indicates a stronger spatial pattern; genes *Foxo1, sst, mog, myl14*, and *ndnf* exhibited clear spatial patterns between polarized estimated hidden indicators. SPARK detected five unique SVGs, as Figure 3(B) shows, for which the PPIs of the estimated hidden expression indicators are close to 0.5. BOOST-HMI identified five unique SVGs, as displayed in Figure 3(C). Among these, seven genes, such as gene *Zfp423, slc5a7* and *palvb*, demonstrated either high or low expression in the majority of cells. The remaining three unique SVGs, *Zic3, Mnat1*, and *slc17a7* delineated three distinct patterns, which may be related to novel biological mechanisms. Lim et al. (2007) previously highlighted the crucial role of *Zic3* in preserving pluripotency in embryonic stem cells, while Herman and El-Hodiri (2002) demonstrated that mutations in *Zic3* are linked to developmental abnormalities such as laterality defects, congenital heart disease, and neural tube defects.

**Figure 3.**
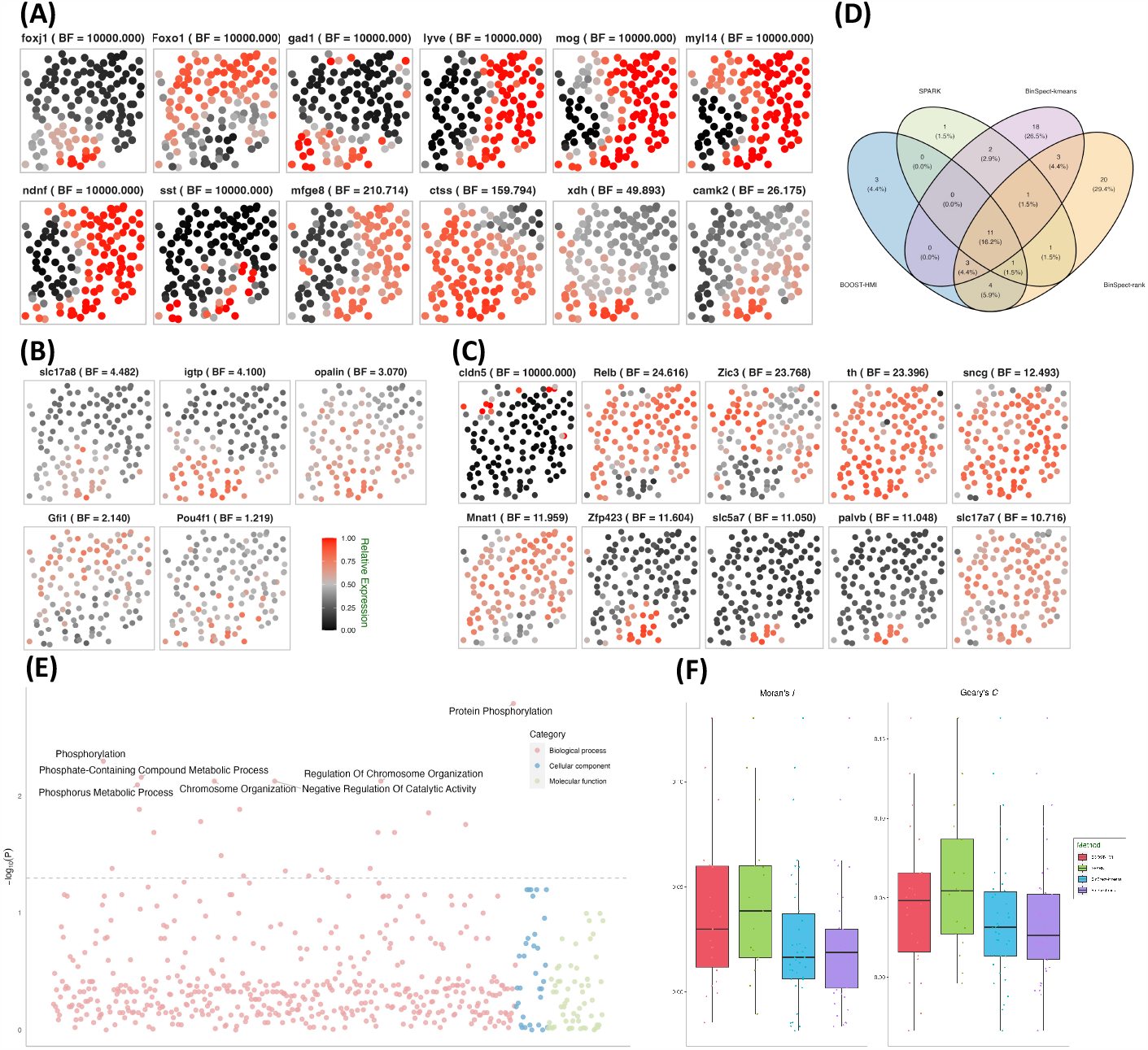
Mouse hippocampus seqFISH data: (A) Spatial distribution of hidden gene expressions ***ξ***^MAP^ of the 12 SVGs detected by both SPARK and BOOST-HMI. (B) Spatial distribution of hidden gene expressions ***ξ***^MAP^ of the five SVGs detected by SPARK only. (C) Spatial distribution of hidden gene expressions ***ξ***^MAP^ of the ten SVGs detected by BOOST-HMI only. (D) Venn diagram of the overlap across SVGs identified by all four methods. (E) Enriched GO terms associated with SVGs detected by BOOST-HMI. (F) Boxplot of Moran’s *I* and Geary’s *C* values for SVGs across the four methods.

In addition to visualizing spatial patterns, we quantified the degree of spatial attraction or repulsion pattern in gene expression across different cells using the spatial autocorrelation tests Moran’s *I* and Geary’s *C*. Moran’s *I* quantifies the spatial clustering or dispersion by standardizing the spatial autocovariance, yielding a correlation coefficient ranging from −1 to 1. A positive Moran’s *I* value corresponds to a spatial clustering pattern where the variable tends to have similar values to its neighboring cells. A Moran’s *I* value close to 0 suggests a random spatial distribution of the data, while a negative value corresponds to a dispersion pattern, where the variable value tends to be dissimilar from its neighbors. To assess the spatial patterns exhibited by the SVGs, Moran’s *I* and Geary’s *C* values were calculated for each SVG identified by at least one of the four methods. Moran’s *I* for each gene *j* was calculated by the following formula:

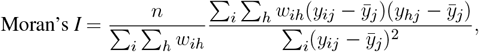

where *w*_*ij*_ = *A/*(*d*_*ih*_)^*m*^ is the connectivity spatial weight between cell *i* and *h*. Spatial weight is a decay factor of the distance between two cells; in our study, we set *A* = 1, *m* = 1. *y*_*ij*_ and *y*_*hj*_ are the gene expression count of cell *i* and cell *h*, and 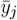*>* is the mean expression of gene *j*. Similar to Moran’s *I*, Geary’s *C* measures the spatial similarity or dissimilarity between neighboring cells, and is calculated with the following formula:

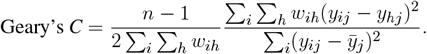

Geary’s *C* ranges from 0 to 2, where a value close to 0 indicates a spatial attraction pattern, 1 corresponds to complete randomness, and 2 implies a spatial repulsion pattern. To ensure uniform interpretation of Moran’s *I* and Geary’s *C*, following Hu et al. (2021), we scaled Geary’s *C* to the range [−1, 1]. The distributions of these values are depicted in Figure 3(F). Remarkably, over 75% of Moran’s *I* and Geary’s *C* values were positive, compellingly indicating the presence of spatial patterns associated with the SVGs across the four methods implemented. Moreover, SVGs from BOOST-HMI and SPARK exhibited the highest Moran’s *I* values, while SVGs from SPARK demonstrated the highest Geary’s *C* values.

To explore the relevant biological functions of identified SVGs, we conducted gene ontology (GO) enrichment analysis using the `R` package `clusterProfiler` (Yu et al., 2012). As mentioned, 214 genes were selected from a list of transcription factors and signaling pathway components. As a result, genes in the background set enriched 4, 622 GO terms and 10, 285 relations. Figure Figure 3(E) depicts the biological processes enriched by SVGs that were detected by BOOST-HMI, such as a smoothened signaling pathway (GO: 0007224), regulation of neural precursor cell proliferation (GO: 20000177), and cellular response to stress (GO: 0033554). Moreover, gene *Foxo1* enriched three significant GO terms, which may inspire further research work on *Foxo1* regulation in the mouse hippocampus. *Mnat1*, one of the SVGs, enriched cellular response to stress. Several studies have found that *Mnat1* is associated with various disease progression and regulation. Qiu et al. (2020) found that *Mnat1*, which was detected only by BOOST-HMI, contributes to the progression of osteosarcoma, and Zou et al. (2020) reported that decreased *Mnat1* expression induces degradation of an important regulator of necroptosis in endothelial cells from samples with Alzheimer’s disease.

### 4.3 Application to the mouse visual cortex STARmap data

The second real dataset we analyzed is a STARmap dataset, which profiles the mouse visual cortex from the hippocampus to the corpus callosum, spanning six neocortical layers at single-cell resolution (Wang et al., 2018). The STARmap dataset measures the expression of 1, 020 genes in 1, 549 cells, including non-neuron cells such as endothelial, oligodendrocytes, astrocytes, and neuron cells, i.e., parvalbumin-expressing, vasoactive intestinal peptide-expressing, and somatostatin-expressing interneurons. Figure 4(A) depicts the layer structure and distribution of cell types within the tissue section as presented in the original study (Wang et al., 2018). Moreover, the STARmap dataset is highly sparse with nearly 79% zero counts. To address potential sources of variability, we performed three quality control steps: 1) cells with fewer than 100 read counts detected were filtered out; 2) genes with more than 90% zero counts were filtered out; 3) genes whose maximum count is smaller than ten were removed. After quality control, the gene expression profile measured the expression of *p* = 107 genes in *n* = 1, 523 cells.

**Figure 4.**
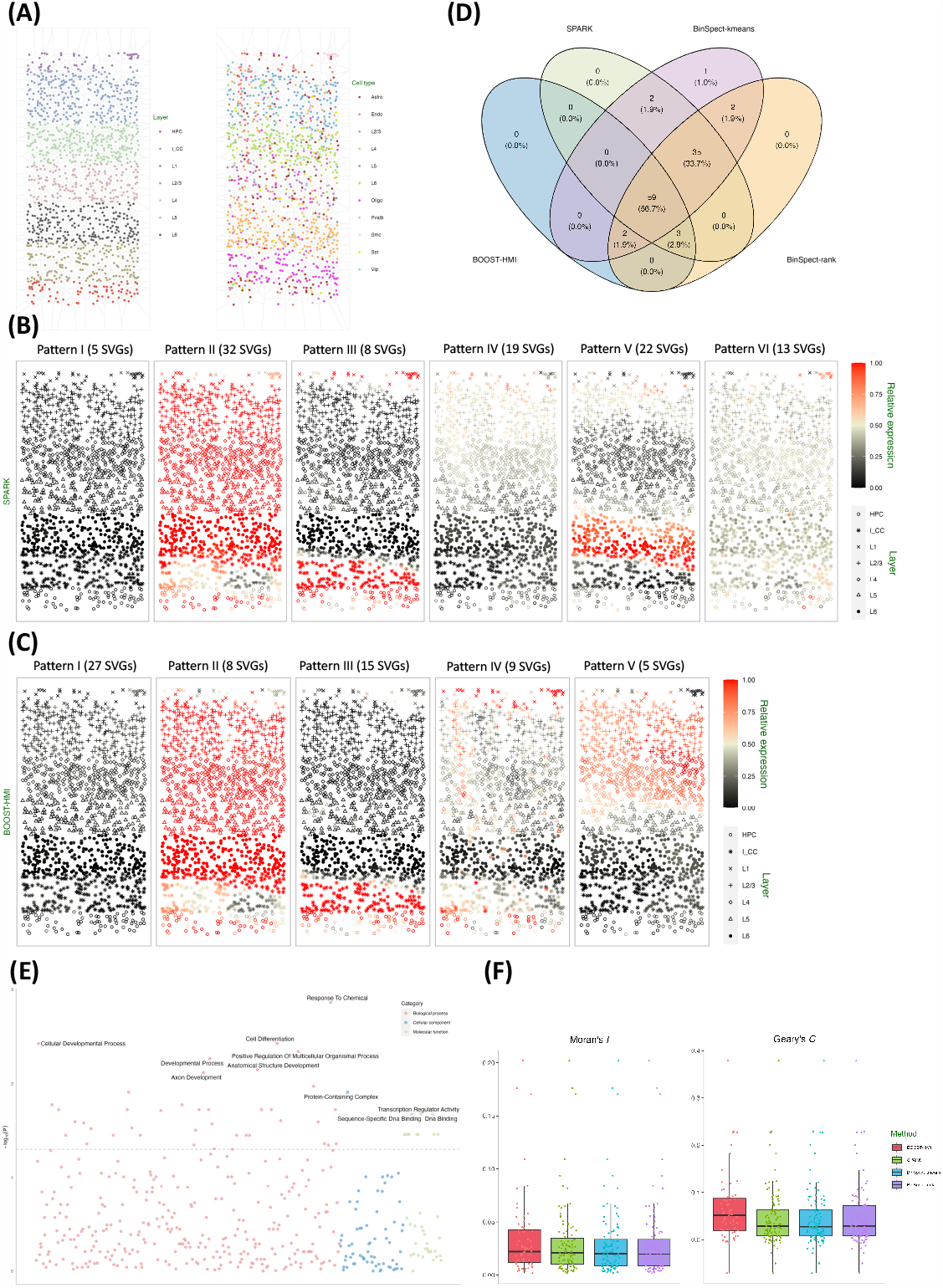
Mouse visual cortex STARmap data: (A) Voronoi diagrams of layer structures and cell type distribution. (B) Spatial distribution of the average hidden gene expressions ***ξ***^MAP^ of the six SVG patterns detected by SPARK. (C) Spatial distribution of the average hidden gene expressions ***ξ***^MAP^ of the five SVG patterns detected by BOOST-HMI. (D) Venn diagram of the overlap across SVGs identified by all four methods. (E) Enriched GO terms associated with SVGs detected by BOOST-HMI. (F) Boxplot of Moran’s *I* and Geary’s *C* values for SVGs across the four methods.

SPARK, BinSpect-kmeans, and BinSpect-rank were implemented with the same parameter settings as the simulation study. As for BOOST-HMI, we set the distance threshold *c*_*d*_ = 0.05 and the decay parameter *λ*_0_ = 60 to satisfy the dependency exp(−*λ*_0_*c*_*d*_) = 0.05. We ran four independent MCMC chains with the same prior specifications and parameter settings as the simulation study, and made posterior inferences after integrating the posterior samples across the four chains.

As Figure 4(D) shows, SPARK detected 99 SVGs, while BOOST-HMI detected 64 SVGs. Both BinSpect-kmeans and BinSpect-rank detected 101 SVGs. SPARK, BinSpect-kmeans, and BinSpect-rank detected 94 SVGs in common. Compared to other methods, BOOST-HMI was conservative, detecting 59 common SVGs with the other three methods. To further investigate the spatial patterns of detected SVGs, we visualized the estimated hidden gene expression indicator for each SVG, annotated with the corresponding Bayes factor, in Figures S3, S4, S5, and S6 of the supplementary materials. Analysis of the detected genes reveals a noteworthy observation: all four methods identify SVGs associated with layer structures, such as *Apod, Apoe*, and *Egr1*, which exhibit high expression in layer L6 and HPC. In contrast, SVGs exclusively detected by the other three methods either lack a clear layer structure or are estimated to be lowly expressed across the entire tissue section, which suggests that BOOST-HMI can detect SVGs with clear spatial patterns and address potential falsification. This conclusion is strongly supported by corroborating evidence from both calculations of Moran’s *I* and Geary’s *C*. Figure 4(F) demonstrates that SVGs detected by BOOST-HMI show stronger spatial autocorrelation than those identified by the other three methods. To delve deeper into the identified spatial patterns, we performed agglomerative hierarchical clustering on the SVGs detected by SPARK and BOOST-HMI, as shown in Figure 4(C). SVGs detected by SPARK were grouped into six clustered, while those detected by BOOST-HMI formed five clusters. Pattern I, II, III and IV from SPARK and BOOST-HMI demonstrate a similar spatial pattern. Spatial pattern V from BOOST-HMI delineates layers L1, L2/3, and L4, while pattern V from SPARK is associated with layer L6. Pattern VI from SPARK highlights a fraction of cells in layer L1.

To gain insights into biological processes, molecular function, and cellular components, GO enrichment analysis was performed on SVGs identified by BOOST-HMI. Figure Figure 4(E) shows that SVGs detected by BOOST-HMI are implicated in biological processes such as the cellular developmental process and anatomical structure development. Additionally, these SVGs are associated with cellular components such as protein-containing complexes as well as molecular functions such as DNA binding. Notably, the gene *Bcl6* significantly enriched the cellular developmental process. Nurieva et al. (2009) found that *Bcl6* functions as a regulator of T follicular helper cell differentiation and B cell-mediated immunity. Our findings have the potential to inspire further novel biological insights.

## 5 CONCLUSION

This paper introduces BOOST-HMI, a novel method for identifying SVGs in imaging-based SRT datasets. By integrating gene expression data with spatial location, BOOST-HMI employs a ZINB mixture model to effectively handle the excessive zeros typical in SRT data. Additionally, it uses a hidden Bayesian mark interaction model to accurately quantify spatial dependencies in gene expressions.

Our approach is adaptable for analyzing sequencing-based SRT data. We validated BOOST-HMI through a simulation study and analysis of two real datasets, demonstrating its effectiveness across various SRT technologies and tissue sections. The simulation results showed that BOOST-HMI is particularly adept at identifying SVGs in data with high sparsity levels, between 30% and 40%. When analyzing the mouse hippocampus seqFISH seqFISH data, BOOST-HMI achieved comparable results to SPARK, with the identified SVGs exhibiting stronger spatial patterns as quantified by Moran’s*I* and Geary’s*C*. Moreover, the SVGs identified were enriched in biologically relevant GO terms, such as smoothened signaling pathways and regulation of neural precursor cell proliferation, offering avenues for further biological investigation.

Further analysis of the mouse visual cortex STARmap dataset revealed that BOOST-HMI can identify SVGs with spatial patterns aligning with the underlying cell structure of the tissue. Additionally, GO enrichment analysis indicated that these SVGs are linked to cellular developmental processes, underscoring the potential for novel biological insights.

While our method assumes homogeneity in spatial patterns across tissue sections, this may not hold true for all cases. Future work will aim to generalize BOOST-HMI to accommodate heterogeneous spatial patterns, enhancing its practicality. Another focus will be on scaling the model to accommodate the growing size of SRT datasets, such as those generated by advanced technologies like Slide-seqV2, which can resolve over 19, 600 genes from around 23, 000 cells (Stickels et al., 2021). Enhanced scalability will enable BOOST-HMI to analyze datasets from various technologies like HDST, Slide-seqV2, and others, potentially leading to more groundbreaking biological discoveries.

## AUTHOR CONTRIBUTIONS

J.Y., X. J., and Q.L. developed the Bayesian framework, designed the MCMC algorithms. J.Y. and X. J. contributed to the review of different methods. J.Y. performed the experiments. X.J., K.W.J., and Q.L. provided resources and helpful discussions. Q.L. and S.S. conceived the study and supervised the application development and the statistical analyses. All authors contributed to the writing of the manuscript. All authors have read and approved the final manuscript.

## FUNDING

This work was supported by the National Science Foundation [2113674,2210912] and the National Institutes of Health [1R01GM141519].

### ACKNOWLEDGMENTS

None.

## SUPPLEMENTAL DATA

The Supplementary Material for this article can be found online at XXX.

## DATA AVAILABILITY STATEMENT

The `R` code and related datasets for this study can be found on GitHub at https://github.com/libedeutch/BOOST-HMI/.

