## Supplementary materials for "Bayesian Hidden Mark Interaction Model for Detecting Spatially Variable Genes in Imaging-Based Spatially Resolved Transcriptomics Data"

### Supplementary Material

#### 1 MCMC ALGORITHM APPENDIX

The model parameter space consists of $(\boldsymbol{H}, \boldsymbol{M}, \boldsymbol{\Phi}, \Xi, \boldsymbol{\omega}_0, \boldsymbol{\theta})$, where $\boldsymbol{H} = \{\eta_{ij}, i = 1, \cdots, n, j = 1, \cdots, p\}$ is the extra zero indicator matrix, $\boldsymbol{M} = \{\mu_{0j}, \mu_{1j}, j = 1, \cdots, p\}$ is the collection of group mean parameters for all genes, $\boldsymbol{\Phi} = \{\phi_{0j}, \phi_{1j}, j = 1, \cdots, p\}$ is the collection of dispersion parameters for all genes, $\Xi = \{\xi_{ij}, i = 1, \cdots, n, j = 1, \cdots, p\}$ is the collection of gene expression level indicator, $\boldsymbol{\omega}_0 = \{\omega_{0j}, j = 1, \cdots, p\}$ is the first-order intensity parameter in the modified energy function, $\boldsymbol{\theta} = \{\theta_j, j = 1, \cdots, p\}$ is the interaction parameter in the modified energy function. The full posterior is,

$$p(\boldsymbol{H}, \boldsymbol{M}, \boldsymbol{\Phi}, \Xi, \boldsymbol{\omega}_0, \boldsymbol{\theta} | Y) \propto f(Y | \boldsymbol{H}, \Xi, \boldsymbol{M}, \boldsymbol{\Phi}) p(\boldsymbol{H}) p(\boldsymbol{M}|\Xi) p(\boldsymbol{\Phi}|\Xi) p(\Xi|\boldsymbol{\omega}_0, \boldsymbol{\theta}) \pi(\boldsymbol{\omega}_0) \pi(\boldsymbol{\theta}).$$

As stated in Section 3 in the main text, the full likelihood and priors are:

$$f(\boldsymbol{Y}|\boldsymbol{H}, \Xi, \boldsymbol{M}, \boldsymbol{\Phi}) = \prod_{i=1}^{n} \prod_{j:\xi_{ij}=0,\eta_{ij}=0} \text{NB}(y_{ij}; s_i\mu_{0j}, \phi_{0j}) \prod_{j:\xi_{ij}=1,\eta_{ij}=0} \text{NB}(y_{ij}; s_i\mu_{1j}, \phi_{1j}),$$

$$\pi(\boldsymbol{H}) = \prod_{i=1}^{n} \prod_{j=1}^{p} \text{Be-Bern}(\eta_{ij}; a_\pi, b_\pi),$$

$$\pi(\boldsymbol{M}|\Xi) = \prod_{i=1}^{n} \prod_{j:\xi_{ij}=0} \text{Ga}(\mu_{0j}; a_\mu, b_\mu) \prod_{j:\xi_{ij}=1} \text{Ga}(\mu_{1j}; a_\mu, b_\mu),$$

$$\pi(\boldsymbol{\Phi}|\Xi) = \prod_{i=1}^{n} \prod_{j:\xi_{ij}=0} \text{Ga}(\phi_{0j}; a_\phi, b_\phi) \prod_{j:\xi_{ij}=1} \text{Ga}(\phi_{1j}; a_\phi, b_\phi),$$

and as described in Section 2.3, the priors in the hidden Bayesian mark interaction model are:

$$\pi(\Xi|\boldsymbol{\omega}_0, \boldsymbol{\theta}) = \prod_{j=1}^{p} \pi(\boldsymbol{\xi}_{\cdot j}|\omega_{0j}, \theta_j),$$

$$\pi(\boldsymbol{\omega}_0) = \prod_{j=1}^{p} \text{N}(\omega_{0j}; \mu_\omega, \tau_\omega^2),$$

$$\pi(\boldsymbol{\theta}) = \prod_{j=1}^{p} \text{N}(\theta_j; \mu_\theta, \tau_\theta^2),$$

where the full formulation of $\pi(\boldsymbol{\xi}_{\cdot j}|\omega_{0j}, \theta_j)$ is shown in Equation (4) in the main text. The p.d.f's of the involved common distributions are given below:

$$\text{If } x \sim \text{NB}(\mu, \phi), \text{ then } p(x) = \frac{\Gamma(x+\phi)}{x!\Gamma(\phi)} \left(\frac{\phi}{\mu+\phi}\right)^{\phi} \left(\frac{\mu}{\mu+\phi}\right)^{x},$$

$$\text{If } x \sim \mathrm{N}(\mu, \sigma^2), \text{ then } p(x) = \frac{1}{\sqrt{2\pi}\sigma} \exp\left(-\frac{(x-\mu)^2}{2\sigma^2}\right),$$

$$\text{If } x \sim \mathrm{Ga}(\alpha, \beta), \text{ then } p(x) = \frac{\beta^\alpha}{\Gamma(\alpha)} x^{\alpha-1} \exp(-\beta x),$$

$$\text{If } x \sim \text{Be-Bern}(a, b), \text{ then } p(x) = \frac{1}{a+b} \frac{\Gamma(a+x)\Gamma(b+1-x)}{\Gamma(a)\Gamma(b)}.$$

Our research interest is to identify SVGs through estimating $\boldsymbol{\theta}$. To estimate $\boldsymbol{\theta}$, all parameters $(\boldsymbol{H}, \boldsymbol{M}, \boldsymbol{\Phi}, \Xi, \boldsymbol{\omega}_0, \boldsymbol{\theta})$ are proposed and estimated in the MCMC algorithm. We use a random walk Metropolis-Hastings (RWMH) algorithm to estimate $\boldsymbol{M}$ and $\boldsymbol{\Phi}$. Due to the intractable normalizing constant in Equation (4), parameters in the hidden Bayesian mark interaction model are estimated by Double Metropolis-Hastings (DMH) algorithm. And $\boldsymbol{H}$ and $\Xi$ are estimated via Gibbs sampler. MCMC algorithm is implemented gene-wisely. The following metropolis-hastings algorithm is described for each gene $j, j = 1, \cdots, p$.

#### 1.1 Random walk Metropolis-Hastings algorithm

**Update of group mean $\boldsymbol{M}$**: We update $\mu_{0j}$ and $\mu_{1j}$ separately, but with the same procedure. Thus, the updating mechanism of $\mu_{0j}$ and $\mu_{1j}$ are integrated as updating $\mu_{kj}$. We propose a new $\mu^*_{kj}$ from $\mathrm{Ga}(a_\mu, b_\mu)$, the proposed $\mu^*_{kj}$ will be accepted with probability $\min(1, r)$. The Hastings ratio $r$ is

$$r = \prod_{i:\eta_{ij}=0, \xi_{ij}=k} \frac{\mathrm{NB}(y_{ij}; s_i\mu^*_{kj}, \phi_{kj})\mathrm{Ga}(\mu^*_{kj}; a_\mu, b_\mu) J(\mu_{kj}; \mu^*_{kj})}{\mathrm{NB}(y_{ij}; s_i\mu_{kj}, \phi_{kj})\mathrm{Ga}(\mu_{kj}; a_\mu, b_\mu) J(\mu^*_{kj}; \mu_{kj})}.$$

Note that the proposal density ratio cancels out for this RWMH update.

**Update of dispersion parameter $\boldsymbol{\Phi}$:** Similar to updating $\boldsymbol{\mu}$, we update $\phi_{0j}$ and $\phi_{1j}$ separately, but with the same procedure. Similarly, the updating mechanism of $\phi_{0j}$ and $\phi_{1j}$ are summarized as updating $\phi_{kj}$. We propose a new $\phi^*_{kj}$ from $\mathrm{Ga}(a_\phi, b_\phi)$, the proposed $\phi^*_{kj}$ will be accepted with probability $\min(1, r)$. The Hastings ratio $r$ is

$$r = \prod_{i:\eta_{ij}=0, \xi_{ij}=k} \frac{\mathrm{NB}(y_{ij}; s_i\mu_{kj}, \phi^*_{kj})\mathrm{Ga}(\phi^*_{kj}; a_\phi, b_\phi) J(\phi_{kj}; \phi^*_{kj})}{\mathrm{NB}(y_{ij}; s_i\mu_{kj}, \phi_{kj})\mathrm{Ga}(\phi_{kj}; a_\phi, b_\phi) J(\phi^*_{kj}; \phi_{kj})}.$$

Note that the proposal density ratio cancels out for this RWMH update.

#### 1.2 Double Metropolis-Hastings algorithm

**Update of first-order intensity parameter $\omega_{0j}$:** We first propose a new $\omega^*_{0j}$ from $\mathrm{N}(\omega_{0j}, \tau^2_\omega)$. We implement the Gibbs sampler to simulate an auxiliary variable $\boldsymbol{\xi}^*_{\cdot j}$ starting from $\boldsymbol{\xi}_{\cdot j}$ based on the new $\omega^*_0$. The proposed value $\omega^*_{0j}$ is accepted to replace the old value with probability $\min(1, r)$. The Hastings ratio $r$ is given as

$$r = \frac{\pi(\boldsymbol{\xi}^*_{\cdot j}|\omega_{0j}, \omega_{1j}, \theta_j)\pi(\boldsymbol{\xi}_{\cdot j}|\omega^*_{0j}, \omega_{1j}, \theta_j)\mathrm{N}(\omega^*_{0j}; \mu_\omega, \tau^2_\omega) J(\omega_{0j}; \omega^*_{0j})}{\pi(\boldsymbol{\xi}_{\cdot j}|\omega_{0j}, \omega_{1j}, \theta_j)\pi(\boldsymbol{\xi}^*_{\cdot j}|\omega^*_{0j}, \omega_{1j}, \theta_j)\mathrm{N}(\omega_{0j}; \mu_\omega, \tau^2_\omega) J(\omega^*_{0j}; \omega_{0j})}.$$

**Update of the interaction parameter $\theta_j$:** We first propose a new $\theta_j^*$ from $\mathrm{N}(\theta_j, \tau_\theta^2)$. We implement the Gibbs sampler to simulate an auxiliary variable $\boldsymbol{\xi}_j^*$ starting from $\boldsymbol{\xi}_j$ based on the new $\theta_j^*$. The proposed value $\theta_j^*$ is accepted to replace the old value with probability $\min(1, r)$. The Hastings ratio $r$ is

$$r = \frac{\pi(\boldsymbol{\xi}_{\cdot j}^*|\omega_{0j}, \omega_{1j}, \theta_j)\pi(\boldsymbol{\xi}_{\cdot j}|\omega_{0j}, \omega_{1j}, \theta_j^*)\mathrm{N}(\theta_j^*; \mu_\theta, \tau_\theta^2)J(\theta_j; \theta_j^*)}{\pi(\boldsymbol{\xi}_{\cdot j}|\omega_{0j}, \omega_{1j}, \theta_j)\pi(\boldsymbol{\xi}_{\cdot j}^*|\omega_{0j}, \omega_{1j}, \theta_j^*)\mathrm{N}(\theta_j; \mu_\theta, \sigma_\theta^2)J(\theta_j^*; \theta_j)}.$$

#### 1.3 Gibbs sampler

**Update of zero-inflation indicator $\boldsymbol{H}$:** The Gibbs sampler is implemented to estimate each $\eta_{ij}, i = 1, \cdots, n$ that corresponds to $y_{ij} = 0$,

$$p(\eta_{ij}|\cdot) \propto \left(\mathrm{NB}(y_{ij}; s_i\mu_{1j}, \phi_{1j})^{1-\eta_{ij}}\right)^{\xi_{ij}} \left(\mathrm{NB}(y_{ij}; s_i\mu_{0j}, \phi_{0j})^{1-\eta_{ij}}\right)^{1-\xi_{ij}} \times \text{Be-Bern}(\eta_{ij}; a_\pi, b_\pi),$$

$$\eta_{ij}|\xi_{ij} = k, \cdot \sim \text{Bern}\left(\frac{\pi(\eta_{ij} = 1|\xi_{ij} = k, \cdot)}{\pi(\eta_{ij} = 1|\xi_{ij} = k, \cdot) + \pi(\eta_{ij} = 0|\xi_{ij} = k, \cdot)}\right).$$

**Update of gene expression level indicator $\boldsymbol{\xi}$:** New $\xi_{ij}$ is generated from $\mathrm{Bern}(p_i)$, where $p_i = \frac{\pi(\xi_{ij}=1)}{\pi(\xi_{ij}=1)+\pi(\xi_{ij}=0)}$. In our implementation, we calculated $p_i$ by $p_i = \frac{1}{\exp(\log(\pi(\xi_{ij}=0))-\log(\pi(\xi_{ij}=1)))+1}$, where $\pi(\xi_{ij} = 1)$ is given in Equation (7) in the main text and $\pi(\xi_{ij} = 0)$ is calculated in the same way.

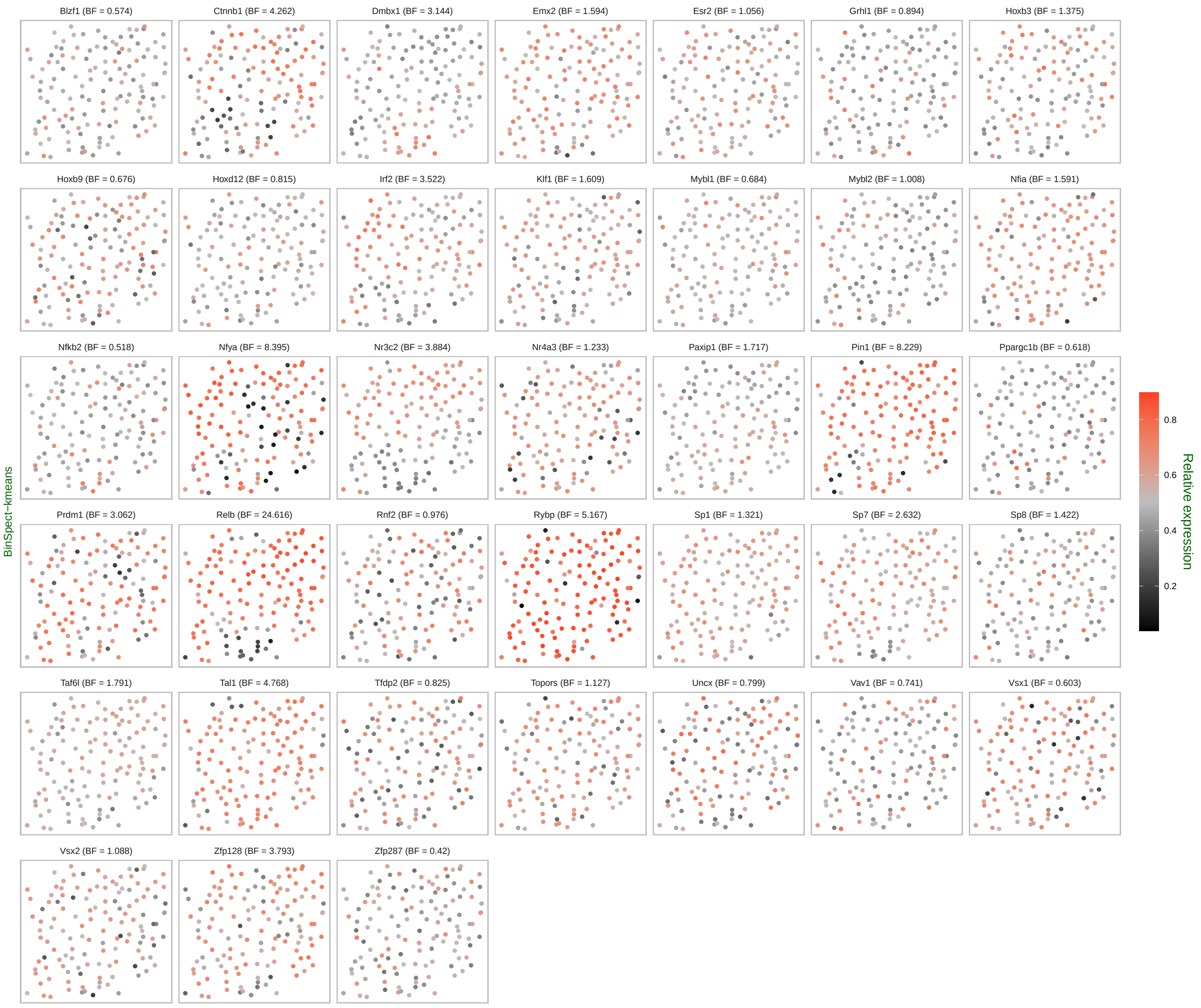


Figure S1: Mouse hippocampus seqFISH data: spatial pattern of hidden gene expression indicator of SVGs detected by BinSpect-kmeans.

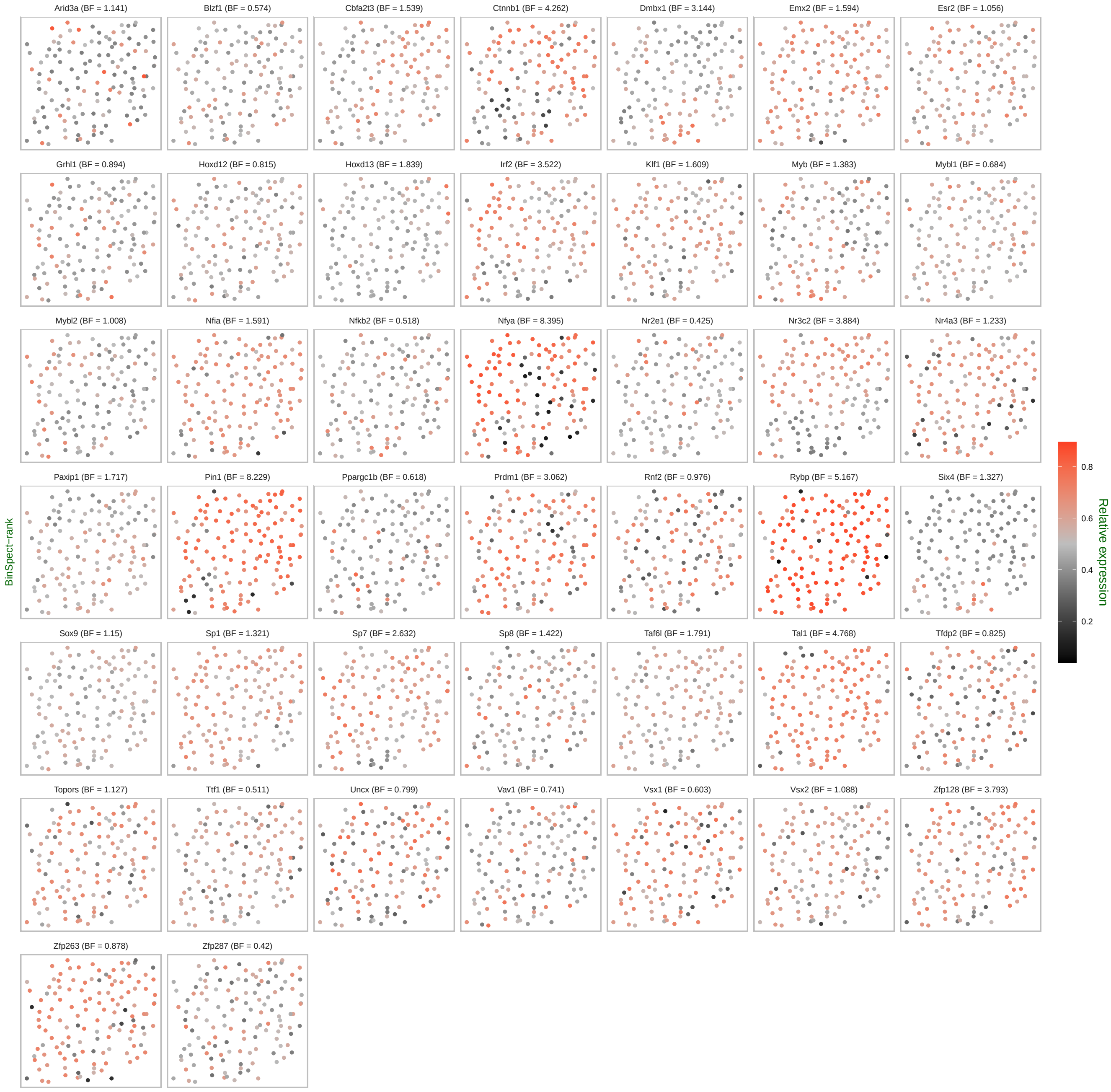


Figure S2: Mouse hippocampus seqFISH data: spatial pattern of hidden gene expression indicator of SVGs detected by BinSpect-rank.

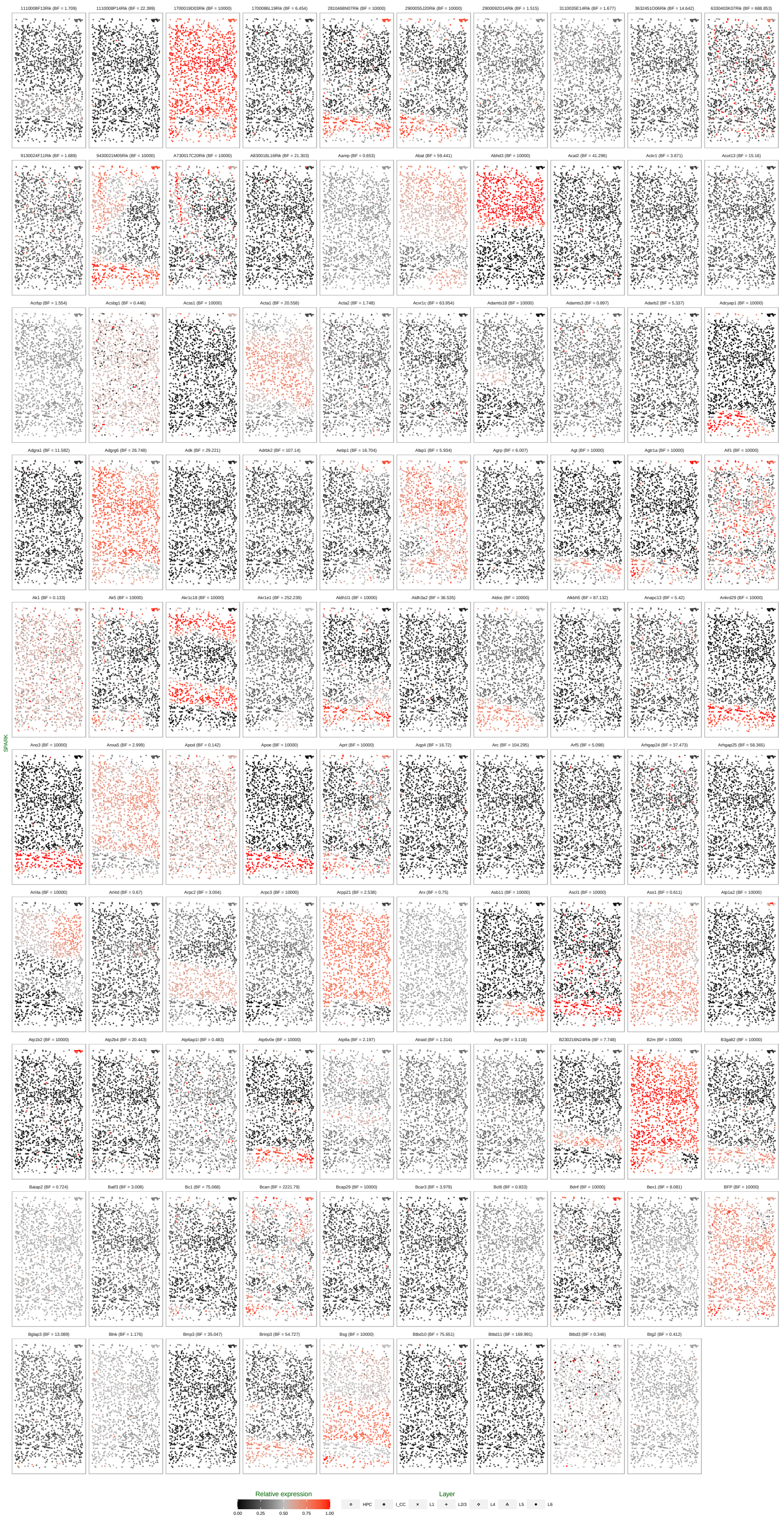

Figure S3: Mouse visual cortex STARmap data: spatial pattern of hidden gene expression indicator of SVGs detected by SPARK.

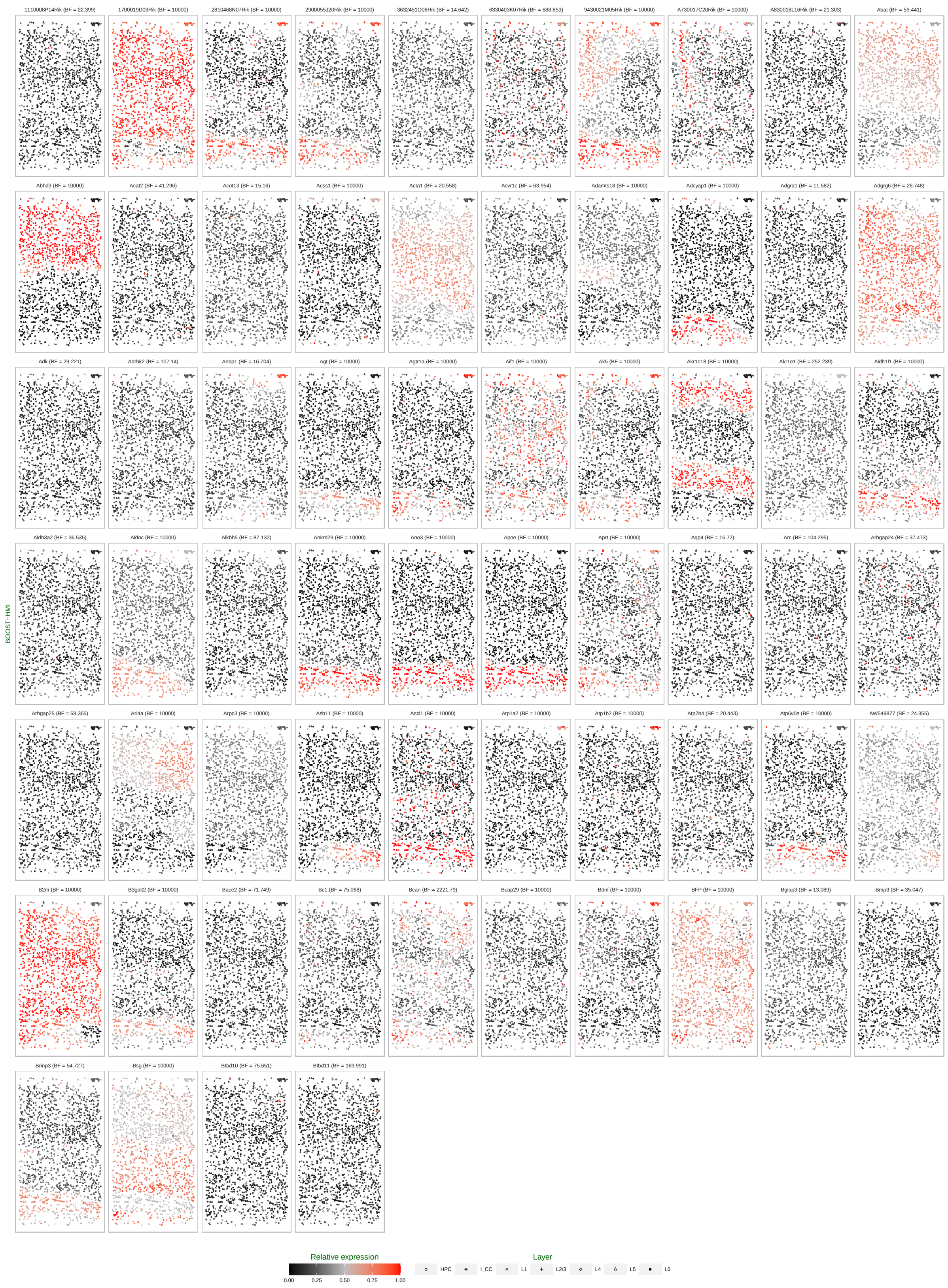


Figure S4: Mouse visual cortex STARmap data: spatial pattern of hidden gene expression indicator of SVGs detected by BOOST-HMI.

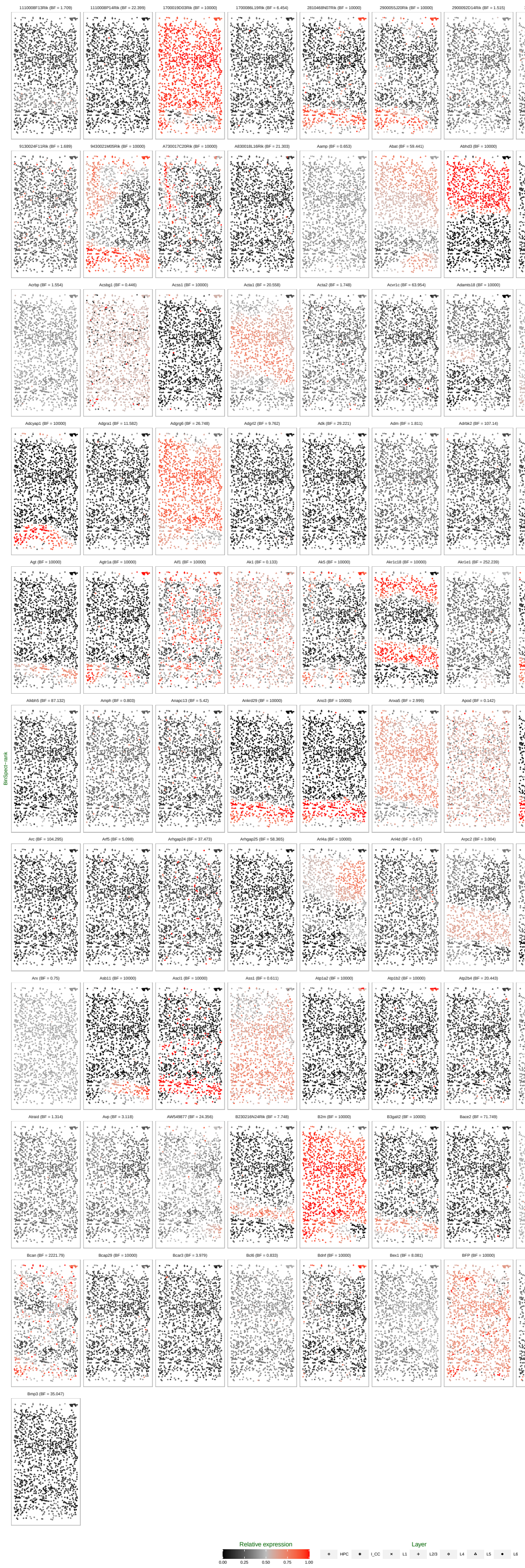

Figure S5: Mouse visual cortex STARmap data: spatial pattern of hidden gene expression indicator of SVGs detected by BinSpect-rank.

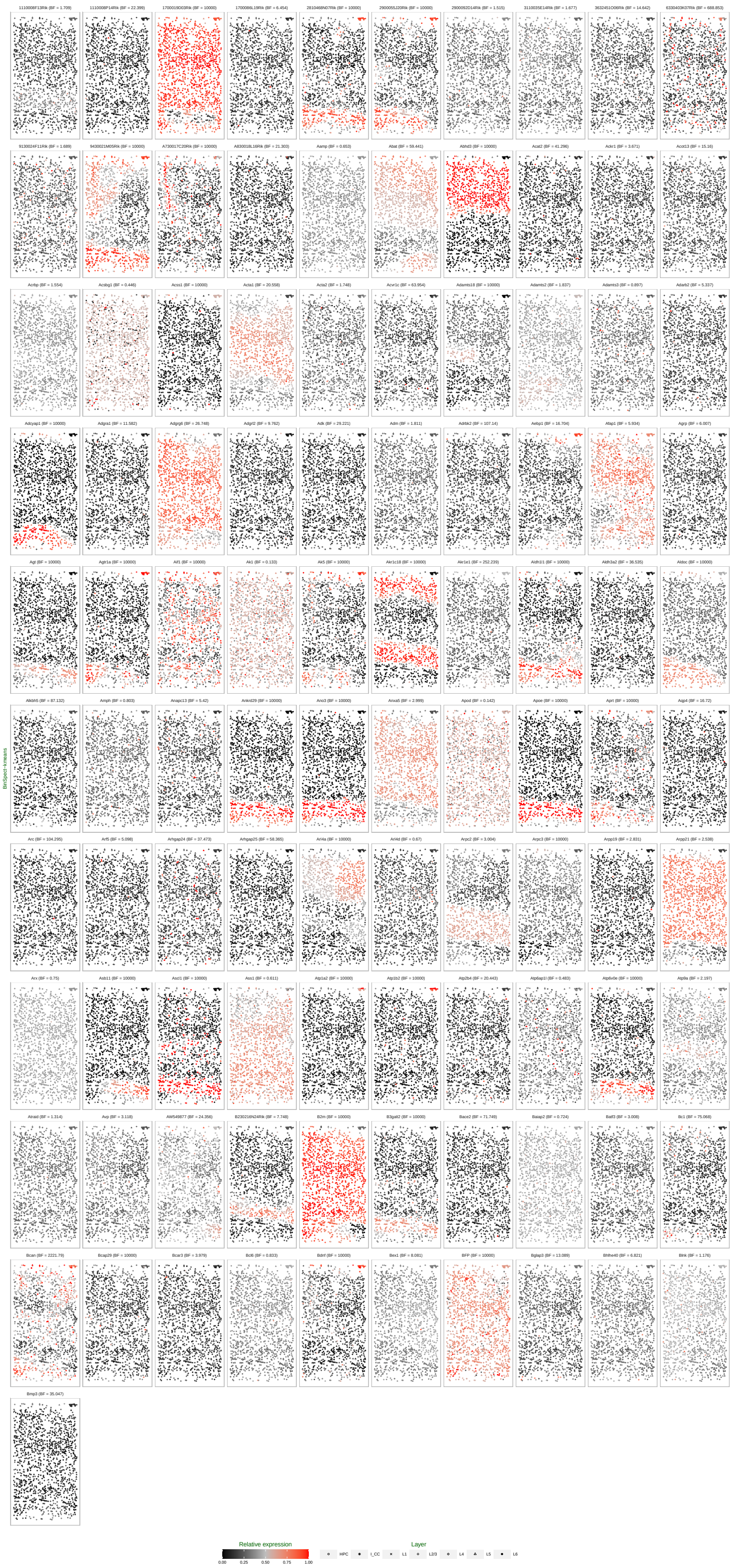


Figure S6: Mouse visual cortex STARmap data: spatial pattern of hidden gene expression indicator of SVGs detected by BinSpect-kmeans.
